# How Autism Impacts Children’s Working Memory for Faces

**DOI:** 10.1101/2025.02.07.636995

**Authors:** Shahrzad M. Esfahan, Narges Sepahi, Ehsan Rezayat

**Affiliations:** Department of Cognitive Sciences, Psychology and Educational Science Faculty, University of Tehran, Tehran, Iran; Institute for Cognitive Science Studies (ICSS), Tehran, Iran; School of Cognitive Sciences, Institute for Research in Fundamental Sciences (IPM), Tehran, Iran

**Keywords:** Working memory, Autism, Face, Children, Precision

## Abstract

This study investigates how visual working memory (WM) functions in children with Autism Spectrum Disorder (ASD) compared to typically developing (TD) children aged 7 to 12, focusing on their ability to remember faces. The primary goal was to understand the differences in visual WM performance between these two groups and identify the sources of memory recall errors. Findings indicate that children with ASD demonstrated significantly poorer visual WM compared to their TD counterparts, particularly in the precision of memory recall. Despite this difference, both groups exhibited similar rates of random guessing, suggesting that the challenges faced by children with ASD are more about accuracy than guessing strategy. This research sheds light on the specific deficits in visual WM associated with ASD, enhancing our understanding of the cognitive mechanisms at play and highlighting potential areas for targeted interventions to support these children.

## Introduction

Autism spectrum disorder (ASD) is a neurodevelopmental condition affecting approximately 23 percent of children (Maenner et al., 2021). It is characterized by deficits in social communication, restricted interests, and repetitive behaviors (American Psychiatric Association, 2013). Various cognitive theories have been proposed to explain the core challenges associated with ASD. Theory of Mind Deficit, Weak Central Coherence, and the Enhanced Perceptual Functioning hypothesis highlight difficulties in social cognition, global information processing, and heightened attention to detail, respectively (Frith, 1989; Happé & Frith, 2006; Mottron et al., 2006). Among these, the Executive Function theory emphasizes deficits in higher-order cognitive processes, including working memory (WM), as a key contributor to ASD-related challenges (Hill, 2004).

WM, a central component of executive function, refers to the ability to temporarily store and manipulate information to guide behavior and cognition (Baddeley & Hitch, 1974). It plays a pivotal role in social communication, enabling individuals to maintain and integrate information during complex interactions (Wang & Gathercole, 2013). Deficits in WM can hinder the ability to process social cues, adapt to changing contexts, and engage in reciprocal communication, all of which are crucial for successful social functioning. Although most studies show deficits in WM of ASD children compared with TD (Gong et al., 2022, Townes et al., 2023; Sadozai et al., 2024), some studies show intact WM in ASD children (Ozonoff et Strayer, 2001; Steele et al., 2007; Macizo et al., 2016, Lynn et al., 2022). Moreover, most studies show deficits in visuospatial WM in comparison with verbal WM (Wang et al., 2017; Habib et al., 2019).

In research on WM in individuals with ASD, Intelligence Quotient (IQ) matching between groups is a commonly used method to control for potential intellectual differences. However, this approach can lead to methodological issues. When IQ is matched, verbal tasks may be subject to underestimation of performance in individuals with ASD, as these individuals often demonstrate specific difficulties with verbal abilities despite having normal or above-average IQ. Conversely, visual-spatial tasks may suffer from overestimation of performance in the ASD group, as individuals with autism may exhibit relative strengths in non-verbal tasks (Russo et al., 2007, Macizo et al., 2016). This can result in misleading conclusions about the cognitive abilities of individuals with ASD. Instead of matching participants based solely on IQ, it is more effective to control for IQ statistically in the analysis. This method allows for a more precise assessment of WM, independent of intellectual ability.

Both delayed match-to-sample and delayed reproduction tasks have been widely used to investigate visuospatial WM in ASD populations. In binary delayed match-to-sample tasks, where responses are limited to “correct” or “incorrect,” studies such as Ozonoff and Strayer (2001) have reported impaired visual memory capacity in individuals with ASD. Alternatively, delayed reproduction tasks involve presenting participants with a stimulus, followed by a delay, and requiring them to identify or reproduce the original stimulus, with responses measured on a graded scale. Using this approach, Maister and Plaisted-Grant (2011) demonstrated deficits in WM among individuals with ASD, whereas Stevenson et al. (2021) observed enhanced precision of visuospatial WM in the same population. These findings highlight the critical influence of task design and stimulus type on WM performance outcomes in ASD.

Social stimuli, such as faces, are particularly relevant in the context of ASD due to the disorder’s hallmark challenges in social cognition. Faces are among the most socially salient stimuli and require the integration of perceptual and memory processes for successful recognition and interaction (Gambarota & Sessa, 2019). Individuals with ASD often show atypical face processing, including reduced attention, lower recognition accuracy, and impaired memory for facial features (Schultz, 2005; Behrmann et al., 2006). These deficits in face processing may further exacerbate difficulties in social communication.

Despite extensive research on non-social stimuli in visual WM tasks, fewer studies have examined visual WM performance for social stimuli in ASD . This study seeks to fill this gap by examining visual WM for faces using a delayed matching-to-sample reproduction task. Data were collected from two groups of children, one with ASD and the other with TD. Our findings reveal that children with ASD exhibit lower precision and higher error rates compared to their TD peers. Including only children allows us to capture the developmental trajectory of visual WM in ASD, which may differ from that in adults (Ozonoff et al., 2001). Additionally, childhood is a key time for interventions aimed at improving social-cognitive abilities, and understanding how visual WM affects these processes can inform more effective strategies (Schultz, 2005). By focusing on children, we ensure that our findings are more relevant to interventions and educational strategies tailored to this age group.

## Method

The present study was a cross-sectional, comparative study between two groups of participants with ASD and TD control group.

### Participants

This study included 20 children diagnosed with ASD and 26 TD children, all aged between 7 and 12 years. Participants had IQ scores above 70 and no evidence of intellectual disability. The TD children had no individual or familial history of ASD or other neurological conditions. The ASD group consisted of children formally diagnosed by a psychiatrist, neurologist, and psychologist. Autism-related behaviors were assessed using the Gilliam Autism Rating Scale, Third Edition (GARS-3). Intelligence quotient (IQ) was evaluated in both groups using the Stanford-Binet Intelligence Scales. Participants were required to have normal or corrected-to-normal vision. Written informed consent was obtained from the parents of all participants prior to the study. The study protocol was approved by the Iran University of Medical Sciences Ethics Committee and adhered to the principles of the Declaration of Helsinki (1964) and its subsequent revisions.

### Stimuli and Procedure

The working memory (WM) test was the same as the one used in Esfahan et al. (2023). Stimuli were displayed on 15.6-inch HD laptop monitors with a 60 Hz refresh rate, with participants seated approximately 40 cm from the screen. During each trial, a face stimulus was presented on the screen, and after variable delays, participants were asked to reproduce the presented face. The stimuli consisted of 11 morphed faces, generated in 10% intervals between two base faces (labeled as Face A and Face B in Figure 1B). These faces were neutral in terms of emotion and gender, with a size of 5° × 6° of visual angle. Stimulus presentation and response collection were implemented using custom routines in MATLAB (MathWorks) with the Psychophysics Toolbox extension 3 (Brainard, 1997; Pelli, 1997; Kleiner et al., 2007).

**Figure 1.**
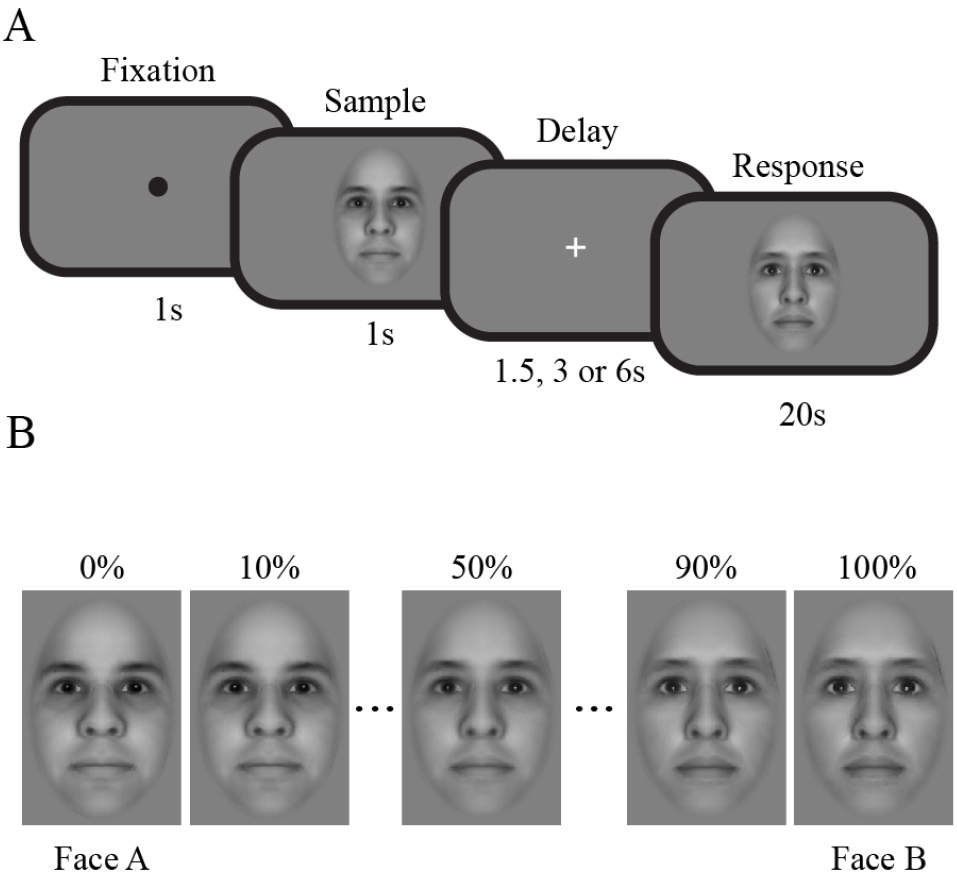
Task overview and procedure. (A) Face working memory task: Each trial included a fixation, ‘sample face’ presentation, a delay, and a ‘test face’ with 20 seconds for response. (B) Morphed faces from face A to face B with 10% interval.

Before the main task, participants received instructions and completed familiarization trials. As shown in Figure 1A, each trial began with a 1-second fixation spot, followed by a 1-second presentation of a “sample face” pseudorandomly selected from four morphed levels of Face A (10%, 40%, 60%, or 90%). After a pseudorandom delay (1.5, 3, or 6 seconds) with a fixation cross, a “test face” was displayed, chosen from one of 10 morph levels excluding the sample face. Participants adjusted the test face using arrow keys to match the sample face and submitted their response with the “SPACE” key. Each block included 36 trials, with participants completing three blocks and optional short breaks between them. Choices and reaction times were recorded.

### Analysis

Following Esfahan et al. (2023), error on each trial was calculated as the difference between the selected “test face” and the “sample face,” ranging from -10 to +10 which were then converted to angular measures from −π to +πradians.

The Mixture Model (Bays et al., 2009) was used to analyze error sources, which include Gaussian variability in memory for the “sample face” and a fixed probability of random guessing.

The maximum likelihood estimates for the probability of reporting the correct answer, the probability of responding randomly, and the κ concentration parameter of the Von Mises distribution were computed separately for each subject and delay. Subsequently, the Von Mises κ was converted into a standard deviation. Precision was determined as the reciprocal of the error’s standard deviation (1/SD), following the approach described by Bays and Husain (2008).

## Results

The study included two groups: 20 children with ASD and 26 TD children. Two participants of ASD and one participant of TD group completed 2 out of 3 blocks. The ASD group comprised 16 males and 4 females, with a median age of 9 years (mean = 9.05, SD = 1.10). The TD group consisted of 14 males and 12 females, with a median age of 9 years (mean = 9.38, SD = 1.55). A Wilcoxon rank-sum test was conducted to compare the ages of the ASD group and the TD group. The results indicated no statistically significant difference in age between the two groups (U=633, z=0.50, p=0.62). The median IQ of the ASD group was 80 (mean = 83, SD = 11.05), and the median IQ of the TD group was 96.5 (mean = 97.58, SD = 11.25). There was a significant difference between the groups (U=791.5, z=4.03, p< 0.0001).

We investigated the root mean square (RMS) error and reaction time in all delays between the ASD and TD groups. The mean RMS error of the ASD group was 1.19 (SD = 0.13), while the mean RMS error of the TD group was 0.92 (SD = 0.17). A Wilcoxon rank-sum test revealed a significant difference between the groups (U=687, z=4.80, p<0.0001), as illustrated in Figure 2A. The mean reaction time of the ASD group was 7.03 (SD = 1.90) and the mean reaction time of the TD group was 6.17 (SD = 1.54), as shown in Figure 2B. Unlike RMS Error, there was no significant difference between the groups (U=547, z=1.70, p=0.09) regarding reaction time.

**Figure 2.**
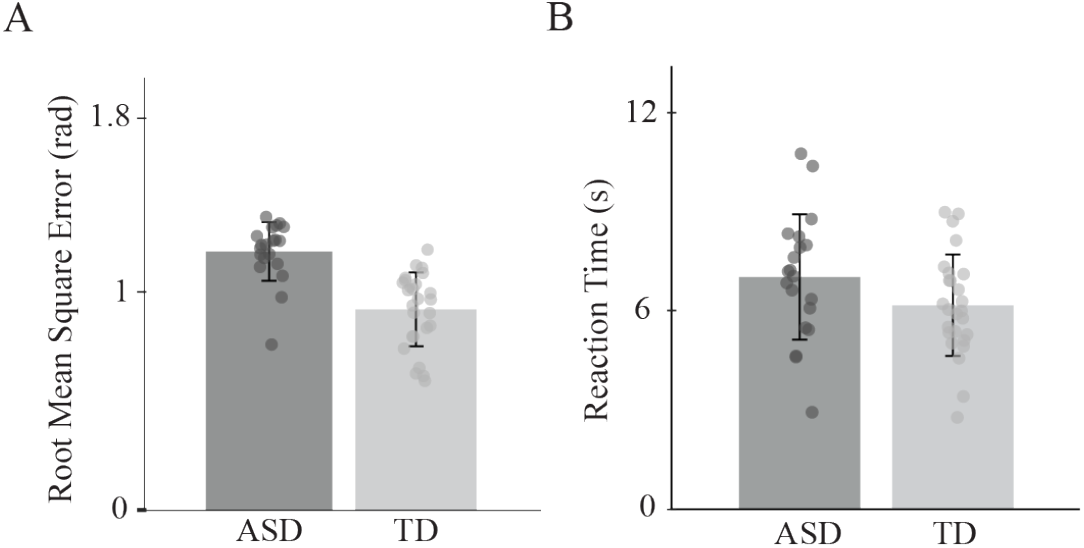
RMS error and reaction time comparisons between ASD and TD groups. (A) Bars represent the mean RMS error for the ASD and TD groups, with error bars indicating the standard error of the mean and circles representing individual RMS error data points. A significant difference in RMS error was observed between the groups. (B) Bars represent the mean reaction time for the ASD and TD groups, with error bars indicating the standard error of the mean and circles representing individual reaction time data points. No significant difference in reaction time was observed between the groups.

We analyzed error across three delay conditions: 1.5, 3, and 6 seconds. At 1.5s delay, there was a significant difference between the ASD and HC groups (U = 431, z = -3.9774, p < 0.001). At 3 s delay, the difference remained significant (U = 420, z = - 4.2212, p < 0.001), and at 6s delay, the difference was also significant (U = 414, z = - 4.3541, p < 0.001).

We utilized the Mixture Model developed by Bays et al. (2009) to examine the distinct sources of error between the two groups. In the context of our task, the model identifies two primary sources of error: (1) Precision error, characterized by the standard deviation (SD) of a Von Mises distribution that represents recall responses centered around the correct answer, and (2) Random guessing, reflected in uniformly distributed errors that are unrelated to the correct answer. For precision error, the results showed that the ASD group had a mean Mixture Model SD of 1.20 (SD = 0.15), while the TD group had a mean Mixture Model SD of 0.88 (SD = 0.21). There was a significant difference between the two groups (U=688, z= 4.82, p < 0.0001, Figure 3A). These findings indicate a greater precision error in the ASD group compared to the TD group. The probability of uniform responses was compared between the ASD and TD groups. The mean probability of uniform responses for the ASD group was 0.0171 (SD = 0.0431), while the mean for the TD group was 0.0230 (SD = 0.0489). A statistical analysis revealed no significant difference between the groups (U=507, z=0.81p = 0.4186, Figure 3B). This suggests that both groups exhibited no significant differences in random guessing.

**Figure 3.**
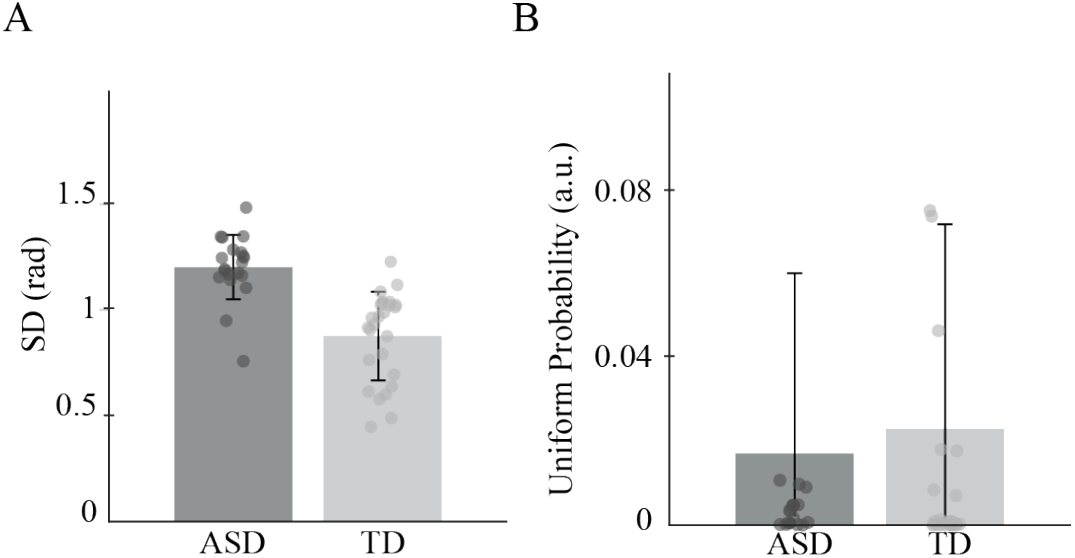
Different sources of error in the face working memory task in ASD and TD group. Precession error and random guess error components were estimated by fitting the mixture model to each participant (see methods). (A) Bars represent the mean of the precession error component for the ASD and TD groups, with error bars indicating the standard error of the mean and circles as the precession error component for each participant. A significant difference in precession error error was observed between the groups. (B) Bars represent the mean of the random guess component for the ASD and TD groups, with error bars indicating the standard error of the mean and circles as the random guess component for each participant. There was not a significant difference between the two groups.

To investigate the effect of delay on RMS error, we conducted a Kruskal-Wallis test across all participants. The results indicated that delay did not significantly impact RMS error overall (H = 1.99, df = 2, p = 0.37). Similarly, no significant effects of delay were observed within the ASD group (H = 4.85, df = 2, p = 0.09) or the TD group (H = 0.83, df = 2, p = 0.66). Additionally, delay had no effect on precision error on RMS error in either the ASD group (H = 2.72, df = 2, p = 0.28) or the TD group (H = 0.82, df = 2, p = 0.66). The probability of random guessing was also unaffected by delay in the ASD group (H = 3.14, df = 2, p = 0.21) or the TD group (H = 1.86, df = 2, p = 0.39). Considering both groups together, neither precision error (H = 0.23, df = 2, p = 0.89) nor the probability of random guessing (H = 4.22, df = 2, p = 0.12) was influenced by delay.

To further investigate the factors contributing to memory errors, an ANCOVA was conducted to examine the effects of group (ASD vs. HC) on mean of RMS error while controlling for IQ. The model included IQ and group as predictors, with error as the dependent variable. The results revealed a significant main effect of group, with individuals in the HC group showing significantly lower RMS error than those in the ASD group (Estimate = -0.23, p < 0.001). However, IQ did not significantly predict error scores (Estimate = -0.002, p = 0.29) when controlling for group membership (R^2^ = 0.44). Continuing to investigate the effects of IQ and group membership on WM performance, an ANCOVA was again performed to investigate the effects of group on the mixture model SD controlling for IQ. The results indicated a significant main effect of group, with the HC group exhibiting lower mixture model SD compared to the ASD group (Estimate = -0.30, p < 0.0001). IQ did not significantly predict SD of precision (Estimate = -0.001, p = 0.57) when controlling for group membership (R^2^ = 0.44). These findings suggest that group membership, but not IQ, plays a critical role in predicting memory errors.

As illustrated in Figure 4, a linear regression was conducted to examine the relationship between IQ and error scores in the ASD and TD groups. In the ASD group, although there was a downward trend indicating a decrease in the mean RMS error with increasing IQ, the results showed that IQ was not a significant predictor of the mean RMS error (R^2^ = 0.15, p = 0.09). Similarly, in the TD group, IQ did not significantly predict the mean RMS error for each participant (R^2^ = 0.17, p = 0.91). However, when the two groups were combined, a significant effect of IQ on RMS error was observed (R^2^ = 0.18, p < 0.01).

**Figure 4.**
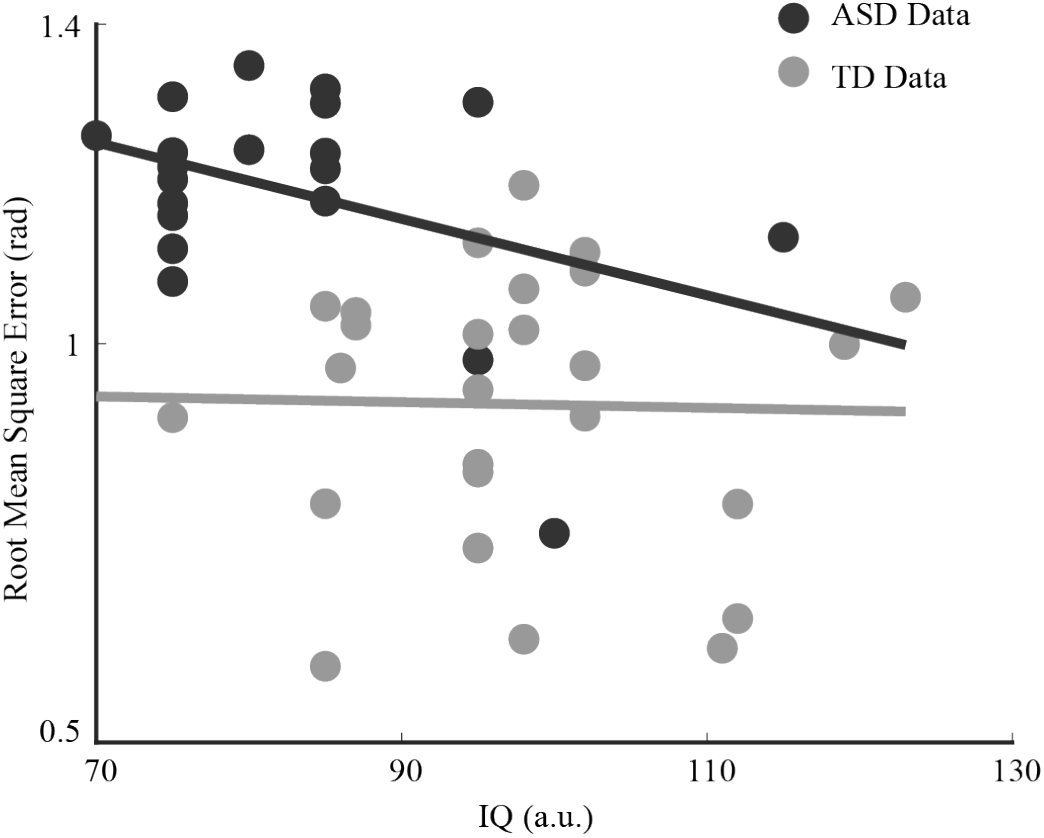
Effect of IQ on mena of RMS error in ASD and TD group. Scatter plot shows individual data points for each participant, with IQ on the x-axis and mean of RMS error on the y-axis. The black and gray lines represent linear regression fits for the ASD and HC groups, respectively. No significant main effect of IQ on error scores was observed in either group. P-values of correlations are provided.

## Discussion

In this study, we observed a significant impairment in visual working memory (WM) performance in children with ASD compared to the TD group. Specifically, we found that precision was significantly lower in the ASD group, while random error did not differ between the groups. These findings align with previous research, which has also reported deficits in visual WM among children with ASD (Wang et al., 2017; Habib et al., 2019; Zhang et al., 2020).

Several studies have demonstrated deficits in WM performance in individuals with ASD compared to TD individuals (Stevenson et al., 2021), primarily using tasks that involve recalling visual features like color and orientation. In this study, we investigated WM using more complex, socially relevant stimuli—faces. Our results revealed that the precision of remembered faces was significantly lower in individuals with ASD. Using faces as stimuli offers distinct advantages. Social stimuli are crucial for understanding ASD, as face processing is a core difficulty for many individuals with the condition. Visual WM for faces is a valuable tool for studying social cognition (Gambarota & Sessa, 2019), providing deeper insights into the social cognitive processes in ASD. Moreover, Given that cortical areas responsible for processing faces differ from those that encode features like color and orientation (Bae et al., 2015), our task explores the role of face-selective areas in WM. Previous research has shown that sensory areas, in addition to frontal areas, play an active role in maintaining information in WM (Ester et al., 2013). Our findings suggest impairments in face-selective cortical areas in ASD, aligning with evidence of abnormal sensory processing patterns in this population (Kuno-Fujita et al., 2020). Moreover, a common limitation of WM tasks using color and orientation is that participants may rely on canonical color categories or cardinal orientations (Pratte et al., 2017). To address this, we used morphed faces that were neutral in terms of gender and emotion, ensuring they were unfamiliar to participants. This design minimizes the potential for biased responses.

While the present study suggests that individuals with ASD show poorer performance on a WM task involving face stimuli, it is important to consider the potential confound of face recognition ability. Previous research has demonstrated that individuals with ASD often struggle with face recognition (Webb et al., 2017). Thus, it is possible that the observed deficit in WM is partially attributable to difficulties in face recognition, rather than an isolated impairment in WM capacity itself. Stevenson et al. (2021) used a delayed-match-to-sample task and applied mixture model to investigate the sources of error. They confound that individuals with ASD exhibited higher precision in WM tasks involving color stimuli, but also showed more frequent binding errors. This suggests that while ASD individuals might have certain advantages in precision, they may also experience challenges with correctly binding or integrating information. The present study, however, used face stimuli. We chose not to involve multiple stimuli in a single trial, as this approach became too difficult for participants. While this eliminated binding errors, it allowed us to focus on precision. We found that precision decreased for the ASD group with face stimuli, suggesting that the social nature of faces introduces challenges that negatively impact WM performance in ASD. Therefore, it is crucial to continue exploring how social and non-social stimuli affect WM in ASD.

In addition to swap errors, participants may also make random errors if items are not properly encoded or retrieved on certain trials, leading to guessing. This source of error, which impairs WM, is represented by random responses. While Stevenson et al. (2021) reported a significant main effect of diagnostic group, with the ASD group exhibiting fewer guesses than the TD group, we found no significant difference in random responses between the ASD and TD groups. Future research should explore this further.

Additionally, response times were not significantly different between groups. This aligns with findings by Ferraro (2016), who reported that while many cognitive and information processing domains are impaired in ASD, simple and choice reaction times appear to remain relatively unaffected.

In our study, the Intelligence quotient (IQ) of children with ASD was lower than that of the control group. However, this variable was controlled for in the analyses, ensuring it does not account for the observed between-group differences. Moreover, participants with intellectual disability (IQ < 70) were excluded, meaning intellectual ability did not influence task performance. Therefore, within the range of average intelligence, our findings demonstrate that IQ does not affect WM performance when comparing ASD and TD groups. This aligns with the findings of Habib et al. (2019), whose meta-analysis similarly concluded that IQ does not explain WM differences in ASD.

Our study had some limitations, including a small sample size and a limited number of tests. Future research should involve larger participant groups and incorporate both social and non-social WM tasks using single and multiple stimuli, as well as additional cognitive assessments. This would provide a more comprehensive understanding of WM in ASD. Additionally, future studies could employ brain imaging techniques, such as functional magnetic resonance imaging (fMRI) or electrophysiological measures, to explore WM performance in relation to the brain areas activated during these tasks.

## Notes

### Competing Interest Statement

The authors have declared no competing interest.

## References

American Psychiatric Association. (2013). Diagnostic and statistical manual of mental disorders (5th ed.). Arlington, VA: American Psychiatric Publishing.

Baddeley, A., & Hitch, G. (1974). Working memory. In G. Bower (Ed.), The psychology of learning and motivation (Vol. 8, pp. 47–89). Academic Press.

Bays, P. M., Catalao, R. F., & Husain, M. (2009). The precision of visual working memory is set by allocation of a shared resource. Journal of Vision, 9(10), 7–7. 10.1167/9.10.7

Bays, P. M., & Husain, M. (2008). Dynamic shifts of limited working memory resources in human vision. Science, 321(5890), 851–854. 10.1126/science.1158023

Behrmann, M., Thomas, C., & Humphreys, K. (2006). Seeing it differently: Visual processing in autism. Trends in Cognitive Sciences, 10(6), 258–264. 10.1016/j.tics.2006.05.001

Brainard, D. H. (1997). The Psychophysics Toolbox. Spatial Vision, 10, 433–436. [PDF]

Ester, E. F., Anderson, D. E., Serences, J. T., & Awh, E. (2013). A neural measure of precision in visual working memory. Journal of Cognitive Neuroscience, 25(5), 754–761. 10.1162/jocn_a_00357

Esfahan, S. M., Nili, M. H. K., Hatami, J., Sanayei, M., & Rezayat, E. (2024). Aging decreases the precision of visual working memory. Neuropsychology, Development, and Cognition B: Aging, Neuropsychology, and Cognition, 31(4), 762–776. 10.1080/13825585.2023.2262105

Ferraro, F. R. (2016). No evidence of reaction time slowing in autism spectrum disorder. Autism, 20(1), 116–122. 10.1177/1362361314559986

Frith, U. (1989). Autism: Explaining the enigma. Oxford: Basil Blackwell.

Gambarota, F., & Sessa, P. (2019). Visual working memory for faces and facial expressions as a useful “tool” for understanding social and affective cognition. Frontiers in Psychology, 10, 2392. 10.3389/fpsyg.2019.02392

Gong, L., Guo, D., Gao, Z., & Wei, K. (2022). Atypical development of social and nonsocial working memory capacity among preschoolers with autism spectrum disorders. Autism Research, 16. 10.1002/aur.2853

Habib, A., Harris, L., Pollick, F., & Melville, C. (2019). A meta-analysis of working memory in individuals with autism spectrum disorders. PLOS ONE, 14(4), e0216198. 10.1371/journal.pone.0216198

Happé, F., & Frith, U. (2006). The weak coherence account: Detail-focused cognitive style in autism spectrum disorders. Journal of Autism and Developmental Disorders, 36(1), 5–25. 10.1007/s10803-005-0039-0

Hill, E. L. (2004). Executive dysfunction in autism. Trends in Cognitive Sciences, 8(1), 26–32. 10.1016/j.tics.2003.11.003

Kleiner, M., Brainard, D., & Pelli, D. (2007). What’s new in Psychtoolbox-3? Perception, 36. [HTML]

Kuno-Fujita, A., Iwabuchi, T., Wakusawa, K., Ito, H., Suzuki, K., Shigetomi, A., Hirotaka, K., Tsujii, M., & Tsuchiya, K. J. (2020). Sensory processing patterns and fusiform activity during face processing in autism spectrum disorder. Autism Research, 13(5), 741–750. 10.1002/aur.2283

Lynn, A., Luna, B., & O’Hearn, K. (2022). Visual working memory performance is intact across development in autism spectrum disorder. Autism Research, 15(5), 881–891. 10.1002/aur.2683

Maenner, M. J., et al. (2021). Prevalence of autism spectrum disorder among children aged 8 years—Autism and Developmental Disabilities Monitoring Network, 11 sites, United States, 2018. MMWR Surveillance Summaries, 70(11), 1–16. 10.15585/mmwr.ss7011a1

Maister, L., & Plaisted-Grant, K. C. (2011). Time perception and its relationship to memory in Autism Spectrum Conditions. Dev Sci, 14(6), 1311–1322. 10.1111/j.1467-7687.2011.01077.x

Mottron, L., Dawson, M., Soulières, I., Hubert, B., & Burack, J. (2006). Enhanced perceptual functioning in autism: An update, and eight principles of autistic perception. Journal of Autism and Developmental Disorders, 36(1), 27–43. 10.1007/s10803-005-0040-7

Ozonoff, S., Pennington, B. F., & Rogers, S. J. (2001). Further evidence of intact working memory in autism. Journal of Autism and Developmental Disorders, 31(3), 257–263. 10.1023/A:1010794902139

Ozonoff, S., & Strayer, D. L. (2001). Further evidence of intact working memory in autism. J Autism Dev Disord, 31(3), 257–263. 10.1023/a:1010794902139

Pelli, D. G. (1997). The VideoToolbox software for visual psychophysics: Transforming numbers into movies. Spatial Vision, 10, 437–442. [PDF]

Pratte, M. S., Park, Y. E., Rademaker, R. L., & Tong, F. (2017). Accounting for stimulus-specific variation in precision reveals a discrete capacity limit in visual working memory. Journal of Experimental Psychology: Human Perception and Performance, 43(1), 6–17. 10.1037/xhp0000302

Russo, N., Flanagan, T., Iarocci, G., Berringer, D., Zelazo, P. D., & Burack, J. A. (2007). Deconstructing executive deficits among persons with autism: Implications for cognitive neuroscience. Brain and Cognition, 65(1), 77–86. 10.1016/j.bandc.2006.04.007

Sadozai, A. K., Sun, C., Demetriou, E. A., Lampit, A., Munro, M., Perry, N., Boulton, K. A., & Guastella, A. J. (2024). Executive function in children with neurodevelopmental conditions: A systematic review and meta-analysis. Nature Human Behaviour, 8(12), 2357–2366. 10.1038/s41562-024-02000-9

Schultz, R. T. (2005). Developmental deficits in social perception in autism: The role of the amygdala and fusiform face area. International Journal of Developmental Neuroscience, 23(2–3), 125–141. 10.1016/j.ijdevneu.2004.12.012

Steele, S. D., Minshew, N. J., Luna, B., & Sweeney, J. A. (2007). Spatial working memory deficits in autism. Journal of Autism and Developmental Disorders, 37(4), 605–612. 10.1007/s10803-006-0202-2

Stevenson, R. A., Ruppel, J., Sun, S. Z., Segers, M., Zapparoli, B. L., Bebko, J. M., Barense, M. D., & Ferber, S. (2021). Visual working memory and sensory processing in autistic children. Scientific Reports, 11(1), 3648. 10.1038/s41598-021-82777-1

Townes, P., Liu, C., Panesar, P., Devoe, D., Lee, S. Y., Taylor, G., Arnold, P. D., Crosbie, J., & Schachar, R. (2023). Do ASD and ADHD have distinct executive function deficits? A systematic review and meta-analysis of direct comparison studies. Journal of Attention Disorders, 27(14), 1571–1582. 10.1177/10870547231190494

Wang, S., & Gathercole, S. E. (2013). Working memory deficits in children with reading difficulties: Memory span and dual task coordination. Journal of Experimental Child Psychology, 115(2), 188–197. 10.1016/j.jecp.2012.11.015

Wang, Y., Zhang, Y. B., Liu, L. L., Cui, J. F., Wang, J., Shum, D. H., & Chan, R. C. (2019). A meta-analysis of working memory in individuals with autism spectrum disorders. PLOS ONE, 14(4), e0216198. 10.1371/journal.pone.0216198

Webb, S. J., Neuhaus, E., & Faja, S. (2017). Face perception and learning in autism spectrum disorders. Quarterly Journal of Experimental Psychology, 70(5), 970–986. 10.1080/17470218.2016.1151059

Zhang, M., Jiao, J., Hu, X., Yang, P., Huang, Y., Situ, M., Guo, K., Cai, J., & Huang, Y. (2020). Exploring the spatial working memory and visual perception in children with autism spectrum disorder and general population with high autism-like traits. PLOS ONE, 15(7), e0235552. 10.1371/journal.pone.0235552

